# Neurodegeneration Associated with Repeated High-Frequency Transcranial Focused Ultrasound

**DOI:** 10.1101/2025.09.23.674044

**Authors:** AP Brna, O Favorov, T Challener, AO Biliroglu, FY Yamaner, RE Kemal, M Annayev, O Oralkan, D Eidum, SB Simons, MP Weisend, PM Connolly

**Author notes:** Corresponding Author: Patrick Connolly.

## Abstract

Transcranial Focused Ultrasound (tFUS) is a popular tool for non-invasive neuromodulation which prior testing paradigms have suggested is benign. However, emerging use cases such as clinical therapies and brain-machine interfaces will likely require repeated or long-duration exposures with novel combinations of stimulus parameters (e.g., frequency, focal volume), and the safety of these conditions has not yet been rigorously validated. Therefore, as an initial study we delivered 1.8 MHz tFUS stimulation to the cortex of 4 non-human primates in 4 sessions each of approximately 90 min over 2 weeks. Motor skills were measured daily with a food pellet picking task. Animals were euthanized and the brain sections were processed for histological markers of neurodegeneration (Fluoro-Jade C). While the animals did not show disruption on the behavioral task, there was clear evidence of neurodegeneration in regions associated with short, intermediate, and extended-duration stimulation, but not with unstimulated control tissues in deeper brain areas. Additional neurodegeneration was observed at locations distant from but functionally connected to stimulated regions, consistent with retrograde damage propagated from neuron processes. We discuss alternative causes for the neurodegeneration, and we recommend the use of additional animal studies to understand this phenomenon, especially for novel stimulation paradigms or parameters and applications where extended or repeated exposures are planned.

## Introduction

Ultrasound is a promising tool for non-invasive neuromodulation with applications in research [1] and the clinic [2]. Low-intensity transcranial focused ultrasound (tFUS) can induce neuromodulatory or neuroplastic changes in neural activity depending on the parameters used [3]. The precise targeting of tFUS makes it attractive both for the treatment of neurological disorders, including Alzheimer’s disease [4] and major depressive disorder [5-7] among many others [8-12], as well as for use in brain-machine interfaces [13-15].

The use of tFUS has to-date been treated as safe. Few indications of negative effects have been observed in animal experiments [16-18], and studies applying tFUS in humans have not revealed any persistent side effects [5, 19, 20]. However, potential therapeutic or neuromodulatory applications of tFUS can require multiple uses over an extended period [5, 6, 21, 22], and few studies have examined the impact of frequent and/or extended tFUS exposure, especially at higher frequencies conducive to fine-grained targeting [23].

In this initial study we examined the effects of extended, repeated exposure to 1.8 MHz tFUS in non-human primates (NHPs). We applied tFUS to each NHP in multiple stimulation sessions and examined targeted cortical sites for signs of neurodegeneration using histological analyses. Despite observing no negative changes in a food pellet picking task, we found neurodegeneration in regions associated with short, intermediate, and extended levels of exposure, as well as at locations distant from but functionally connected to stimulated regions. Our results raise concerns on the safety of tFUS in the tested regime and suggest the need for additional studies to examine longer-term effects of tFUS prior to routine use.

## Methods

### NHP Exposure Study

We exposed 4 squirrel monkeys (1 male, 3 female) to tFUS stimulation and measured the effects on cortical tissue and behavior. We ran 4 stimulation sessions on each NHP over a 2-week period. Behavioral performance on the food pellet-picking task was monitored for at least 19 days prior to any procedures and for at least 3 days following the final stimulation session. Next, we ran histological analyses on the cortex in stimulated regions to test for neurodegeneration.

Each NHP underwent 2 stimulation sessions per week on separate days for 2 weeks for a total of 4 sessions. Sessions within the week were either consecutive or 1 day apart, though each NHP experienced at least 1 pair of consecutive sessions. Sessions consisted of 2 hours of stimulation in the sensorimotor region of one hemisphere, which was split between a single extended-duration site and 1 or more short or intermediate-duration sites. Figure 1e illustrates the pulse train repeats we applied to a given site using terminology defined by [24]. We chose parameters to induce neuron excitation [25, 26] using intensities effective at higher operating frequencies (*f*_*0*_) [27]. Intensities were determined through individualized simulations of ultrasound propagation using Fullwave software [28] (Figure 1d) (see tFUS Hardware and Targeting), and we calculated the total exposure time (*T*_*E*_) of each site as the cumulative duration of all active tFUS pulses applied there (Figure 1e). Intensity and duration details are listed in Table 1 and plotted in Figure 2.

**Table 1.** Stimulation site details. Intensities calculated from pressures simulated in Fullwave as described in text, and focal volume represents -3dB drop from maximum intensity at each site. *TE* calculated from pulse train duty cycle (*DC*_*Train*_) = 50% and cumulative pulse trains applied to each site.

| NHP | Site Type | Derated ISPPA (W/cm<sup>2</sup>) | Derated ISPTA<sub>Train</sub> (W/cm<sup>2</sup>) | Exposure Time $T_E$ (sec) | Focal Volume (mm<sup>3</sup>) |
| --- | --- | --- | --- | --- | --- |
| A | Extended | 37.5 | 18.8 | 119.8 | 1.7 |
|  | Intermediate | 35.6 | 17.8 | 25.5 | 1.6 |
|  | Short | 37.2-44.8 | 18.6-22.4 | 3.0-6.4 | 1.5-1.8 |
| B | Extended | 22.7 | 11.3 | 132.0 | 0.4 |
|  | Short | 35.1-65.4 | 17.5-32.7 | 4.5-6.8 | 0.3-2.2 |
| C | Extended | 18.2 | 9.1 | 133.2 | 3.3 |
|  | Intermediate | 38.5 | 19.2 | 40.8 | 3.0 |
| D | Extended | 23.8 | 11.9 | 133.2 | 3.4 |
|  | Intermediate | 44.9 | 22.4 | 40.8 | 3.3 |

**Figure 1.**
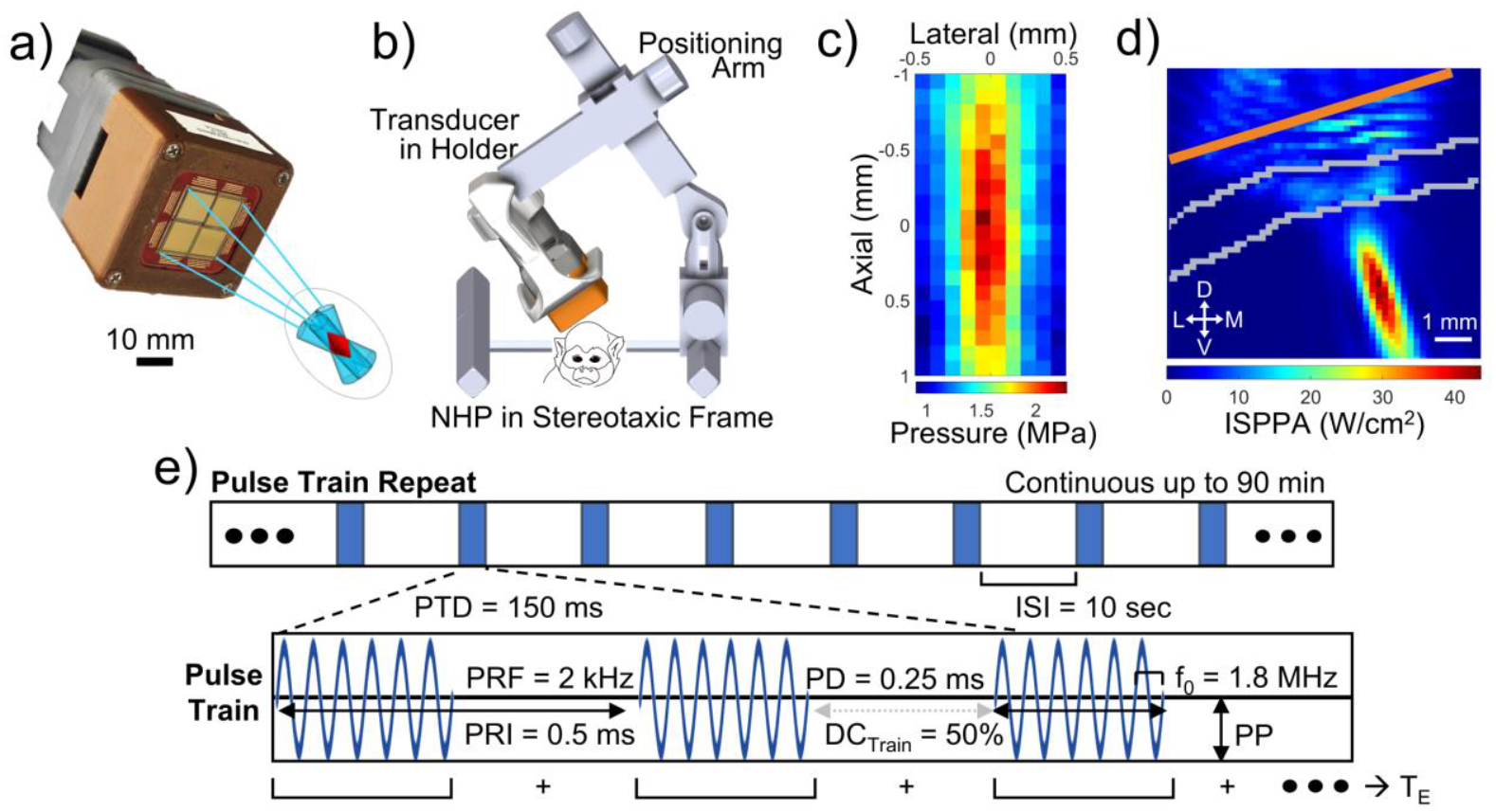
Stimulation setup. (a) Phased array transducer used for tFUS stimulation. Delays loaded into 1-4 arrays allow focusing and steering of sound waves for neuromodulation. Only 1 array was used in this work. (b) Experimental setup showing NHP and transducer placement within stereotaxic frame. (c) Profile of ultrasound focus in free field captured with a hydrophone. Image color denotes peak ultrasound pressure amplitude. (d) Fullwave simulation of ultrasound focus for short-duration site in NHP A. Orange line denotes ultrasound array placement, and skull location is outlined in gray. Image color denotes ultrasound spatial-peak pulsed average intensity (*ISPPA*) with 1 mm scale bar and directional compass (M = medial, L = lateral, D = Dorsal, V = Ventral), at bottom. (e) Description of applied tFUS pulse train repeats. Each pulse train repeat consisted of multiple pulse trains, where each pulse train was applied to a single site (extended or intermediate-duration) or to multiple sites in sequence (short-duration). *PTD* = pulse train duration, *ISI* = inter-stimulus interval, *PRF* = pulse repetition frequency, *PRI* = pulse repetition interval, *PD* = pulse duration, *DC*_*Train*_ = duty cycle of pulse train, *f*_*0*_ = operating frequency. *PP* = peak pressure. *T*_*F*_ = exposure time. Terminology follows [24].

**Figure 2.**
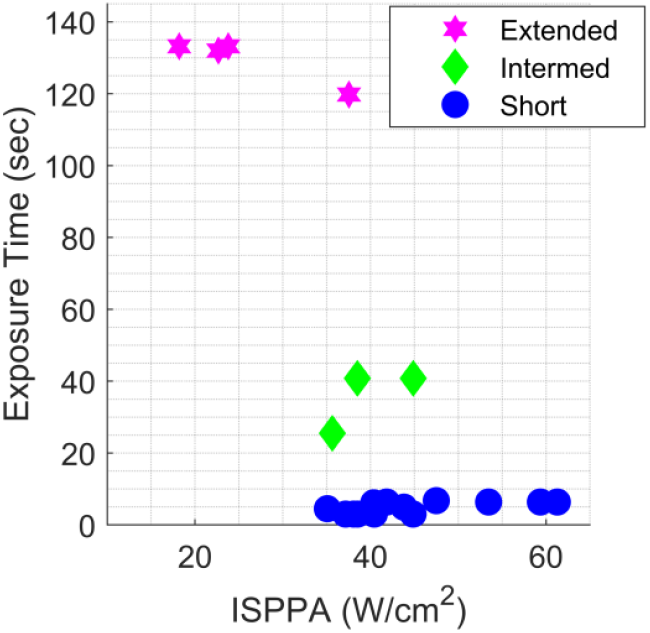
Stimulation site exposure details. *ISPPA* and *TE* for each stimulation site across all NHPs, with site types denoted by color/shape. Broadly, sites with higher intensities were exposed to tFUS for shorter durations in total, and sites stimulated for longer durations tended to be lower intensity.

In the first 2 NHPs (A, B), we applied tFUS to extended-duration and short-duration sites and a 1-day intermediate-duration site. The extended-duration site for each NHP was placed anterior of the central sulcus in motor cortex (Figure 3a-b); we applied pulse train repeats to these sites on average 81.1 min per session. 7 short-duration sites were arranged as a line running mediolaterally and centered approximately 2 mm anterior and 2 mm lateral from the extended-duration site; sites were spaced every 400 μm. We applied pulse train repeats to each individual site for less than 11 min per session. In NHP A, we also applied pulse train repeats to an intermediate-duration site 2.4 mm posterior of the extended-duration site for 1-day for 57.5 min.

**Figure 3.**
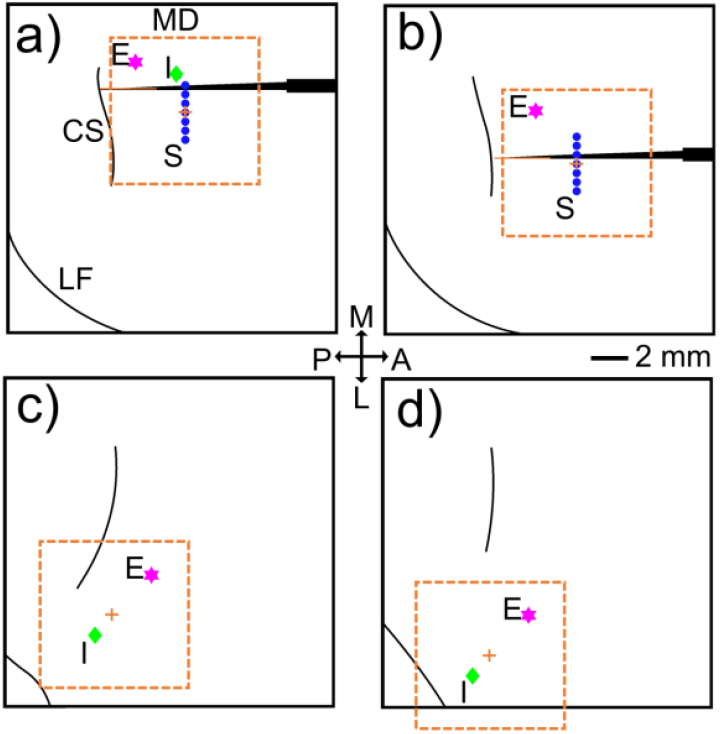
Stimulation site locations. Markers Blue dots give locations for stimulation sites on the sensorimotor cortex of each NHP. Magenta stars are extended-duration sites also marked (E), green diamonds are intermediate-duration sites also marked (I), and blue circles are short-duration sites also marked (S). Individual NHPs A-D shown on corresponding panels (a-d). Panels are rotated and scaled to a common orientation as denoted by the 2 mm scale bar, directional compass (M = medial, L = lateral, A = Anterior, P = Posterior), and cortical landmarks (MD = midline along panel top, CS = central sulcus, LF = lateral fissure). Orange boxes and plus signs show size, shape, and center of active tFUS array relative to stimulated locations. NHPs A and B also show inserted probes in black, with recording electrode region at tip shown in different color. No stimulated sites were within 1 mm of recording electrode locations.

Additionally, we inserted a single-shank chronic recording probe (NeuroNexus) tangentially into the sensorimotor region of both NHP A and B to record single unit responses near the short-duration sites. Stimulation sessions were run at least 1 week after electrode probe insertion for recovery. We observed during slice preparation that the recording electrode placements were far (>1 mm) from stimulated sites (Figure 3a-b), precluding analysis of stimulation responses.

To avoid histological confounds, we did not implant electrodes nor perform surgery in the second 2 NHPs (C, D). We applied tFUS to a single extended-duration and a single intermediate-duration site each stimulation session. The extended-duration site for these NHPs was placed near to the central sulcus, targeting sensorimotor regions associated with the hand. The intermediate-duration site was placed approximately 3 mm posterior and 3.5 mm lateral from the extended duration site in each NHP (Figure 3c-d). We applied pulse train repeats to the extended and intermediate-duration sites for 75.1 and 23.0 min per session, respectively.

### tFUS Hardware and Targeting

We used an electronically-steered ultrasound transducer to deliver tFUS [23]. The stimulator (Figure 1a) uses capacitive micromachine ultrasonic transducers (CMUTs) in a 2D phased array to steer and focus ultrasound using constructive interference [23, 29]. The transducer array consisted of 1024 CMUT elements arranged in an 8x8 mm square bonded to a custom application-specific integrated circuit (ASIC)to control the phase and amplitude of ultrasound output from each element. Using an *f*_*0*_ of 1.8 MHz, this transducer produced sinusoidal pressure waves at -3 dB focal volumes of 0.2-3.4 mm^3^, depending on the target site.

We calculated transducer phase delays and derated peak pressures for each target site using finite-difference methods for nonlinear ultrasound propagation using Fullwave software (University of North Carolina at Chapel Hill) [28]. Initial target sites were chosen by co-registering CT scans of each NHP’s skull with a squirrel monkey MRI atlas [30] and identifying regions corresponding to sensorimotor representations of the contralateral, dominant hand. NHP scans, target sites, and transducer placement information were merged in Fullwave to visualize focal regions and calculate optimal phase delays for each transducer element given individual skull morphologies (Figure 1d). Focal regions were ovoid from the center of the transducer to target sites with -3 dB lateral and axial focal widths of approximately 400 μm and 2 mm, respectively. Longer-duration sites were placed farther from the central axis of the transducer, making their focal regions comparatively more transverse. Target depths were 2-2.75 mm beneath the skull to avoid focusing in bone.

Additionally, we verified our targeting method, volumes, and pressure distributions with *ex vivo* measurements of tFUS transmission through NHP-B’s excised skull using a hydrophone in a water bath after the completion of its 4 stimulation sessions. The peak pressure amplitude produced under free-field conditions focused 7.5 mm beneath the center of the transducer array mimicking a short-duration site was measured at 2.3 MPa (Figure 1c). Volume and pressure profiles under both free-field and through-skull conditions were comparable to Fullwave simulations.

### Metric Calculations

We calculated intensities using Equations 1 and 2 from derated peak pressure amplitudes derived from Fullwave simulations at each target site.

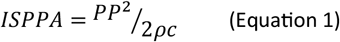

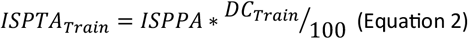

where *ISPPA* is spatial-peak pulsed average intensity (W/m^2^), *ISPTA*_*Train*_ is spatial-peak temporal average intensity for pulse train (W/m^2^), *PP* is the peak pressure amplitude (Pa), *ρ* is cortical tissue density (1000 kg/m^2^), *c* is the speed of sound in cortical tissue (1468.4 m/s), and *DC*_*Train*_ is the duty cycle of a pulse train (%).

As shown in Table 1, *ISPPA* values varied across target sites, but broadly extended-duration sites were lower-intensity, intermediate-duration sites were intermediate intensity, and short-duration sites were higher-intensity. The largest *ISPPA* used was 65.4 W/cm^2^ at a short-duration site. We calculated *ISPTA*_*Train*_ using *DC*_*Train*_ to reflect potential for heating over the course of a single stimulus, though other conventions incorporating *ISI* (*ISPTA*_*Train-Repeat*_) are also commonly used [16, 24].

We calculated temperature change in cortical tissue (*ΔT*) and the mechanical index (*MI*) to estimate thermal and mechanical impacts. Equations used for *ΔT* are given in Equations 3-4, which were taken from [27] and adjusted for use with a pulsed waveform assuming no temperature loss during off-periods.

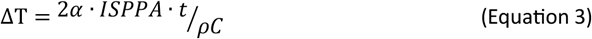

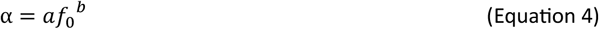

where *ΔT* is the change in temperature (°C), *t* is *PTD* · *DC*_*Train*_/100 (sec), *C* is the specific heat of cortical tissue (3640 J/kg/K), *f*_0_ is the operating frequency (MHz), *a* is the absorption coefficient of cortical tissue (2.4 Np/m/MHz^-b^), and *b* is the power law coefficient (1.18).

The equation used to calculate *MI* is given in Equation 5:

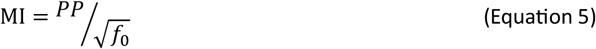

where *MI* is the mechanical index, *PP* is the negative peak pressure amplitude (MPa), and *f*_0_ is the operating frequency (MHz). We calculated our maximum *ΔT* and *MI* in cortical tissue to be 0.13°C and 1.0, respectively. The largest value for each metric also corresponded to the highest-intensity short-duration site.

### Stimulation Session Procedure

Anesthesia was induced with 4% isoflurane. After intubation, the NHP’s head was shaved then secured within a stereotaxic frame (Figure 1b). Head position and tilt were set using 2 frame-mounted bars inserted into the ears and a hooked bar at the lowest point of an orbital socket. The tFUS transducer was mounted to a positioning arm using a 3D-printed holder and was placed at the precalculated position, 1-2 mm from the scalp. Markings applied to the scalp verified placement across multiple sessions. Ultrasound gel centrifuged to remove any bubbles coupled the transducer to the scalp. During stimulation, light anesthesia was maintained using 0.4-1.5% isoflurane with a 50%/50% oxygen and nitrous oxide gas mixture. Heart rate, rectal temperature, and breathing rate were monitored at regular intervals with breathing rate controlled through a ventilator. All stimulation sessions, behavioral tests, and surgeries were monitored by veterinary staff and carried out according to protocols approved by the Institutional Animal Care and Use Committee of the University of North Carolina at Chapel Hill.

Stimulation was applied in pulse train repeats [24] with 150 ms pulse trains with a 10 second *ISI* for up to 90 min (Figure 1e). In NHPs A and B, we stimulated the 7 short-duration sites with 20 pulse train repeats applying a single pulse train to each site in sequence. Afterwards, for 1 or more individual sites we applied up to 45 pulse trains to investigate tFUS-driven changes in activity. Finally, we applied pulse train repeats to other sites until 2 hours had elapsed following anesthesia induction. In NHP A, we stimulated the intermediate-duration site in the first session and the extended-duration site for remaining 3, and in NHP B we stimulated the extended-duration site for all 4 sessions. For each session in NHPs C and D, we applied 75.1 minutes of pulse train repeats exclusively targeting the extended-duration site, approximating the average *T*_*E*_ at extended-duration sites of NHPs A and B. We immediately followed this with 23.0 min of pulse train repeats exclusively targeting the intermediate-duration site. Given our low *ΔT* and *MI* values, we did not expect neurodegeneration from such stimuli.

### Behavioral Assessment

As an additional measure of cortical health, we used a food pellet picking task to evaluate if any damage observed in histology correlated with decrements in behavioral performance. We trained each NHP to pick food pellets from a rotating table, which requires gross and fine motor skills and sensory feedback to accomplish. Task performance was measured prior to, during, and after the completion of all 4 stimulation sessions. The rotation speed of the table was adjusted for each NHP to achieve approximately an 80% success rate prior to any stimulation sessions then held constant through the study.

### Histology

After the completion of their last stimulation session, each NHP was euthanized with 5% isoflurane and perfused. Cortical tissue was collected and fixed for subsequent analysis. After brain extraction, 50 μm sagittal sections were made from regions surrounding tFUS targets, and these were subsequently stained with Fluoro-Jade C (FJC) and DAPI. FJC is an indicator for neuron degeneration regardless of mechanism [31], and DAPI reveals cell nuclei [32].

Cortical sections were stained using the free-floating method. Briefly, sections were permeabilized in 0.3% Tris Triton for one hour at room temperature, then blocked using 10% normal goat serum in 0.3% Tris Triton for one hour at room temperature. Sections were incubated in primary antibodies for one stain type [results not shown, under review] for 48 hours at 4°C, rinsed, then incubated in secondary antibodies for that same stain for three hours at room temperature. Sections were rinsed again, mounted onto charged slides using water gelatin, and dried overnight at room temperature. After rehydration, sections were stained with FJC and DAPI according to the kit instructions (Biosensis, #TR-100-FJ). The incubation time for FJC plus DAPI was increased to 1.25 hours. Finally, slides were dehydrated at 60°C for 15 minutes, cleared briefly in xylene, and coverslipped using DPX (Electron Microscopy Sciences, #13512).

For quantitative comparison of FJC labeling at different tFUS-stimulated cortical sites, at each such site a 1.25×1.25 mm box was placed over the most densely labeled region in the most densely labeled cortical section. The distance between every pair of the labeled cells in the box was computed and the shortest such distance was identified for every labeled cell in the box. The FJC labeling density was then computed as the mean shortest distance in the box (MSD). Lower MSD values reflect higher labeling density. Statistical significance of the difference in MSD values between two stimulated sites was computed with Welch’s t-test, using 50 labeled cells sampled at random in each site.

## Results

### FJC Labeling at tFUS-targeted Sites

We observed FJC-labeled cells at cortical locations targeted by tFUS-stimulation in all 4 NHPs (Figures 4 and 5). In many labeled cells, FJC can be seen filling triangular-shaped bodies and proximal apical dendrites, making it clear that these cells are pyramidal neurons, but other labeled cell profiles were more circular or polygonal and without visible processes, making it hard to characterize them (Figure 5f-g). Some of such uncategorized cells could be inhibitory interneurons or glial cells [33]. In NHPs A and B, pyramidal cells were most evident in cortical layers 2-3 but were also very sparsely distributed in deeper layers 5-6; uncategorized cells were spread more broadly and across all cortical layers, but still at greater density in the upper layers (Figure 5a-c). In contrast, both pyramidal and uncategorized cells were restricted almost exclusively to layers 2-3 in NHPs C and D (Figure 5d-e).

**Figure 4.**
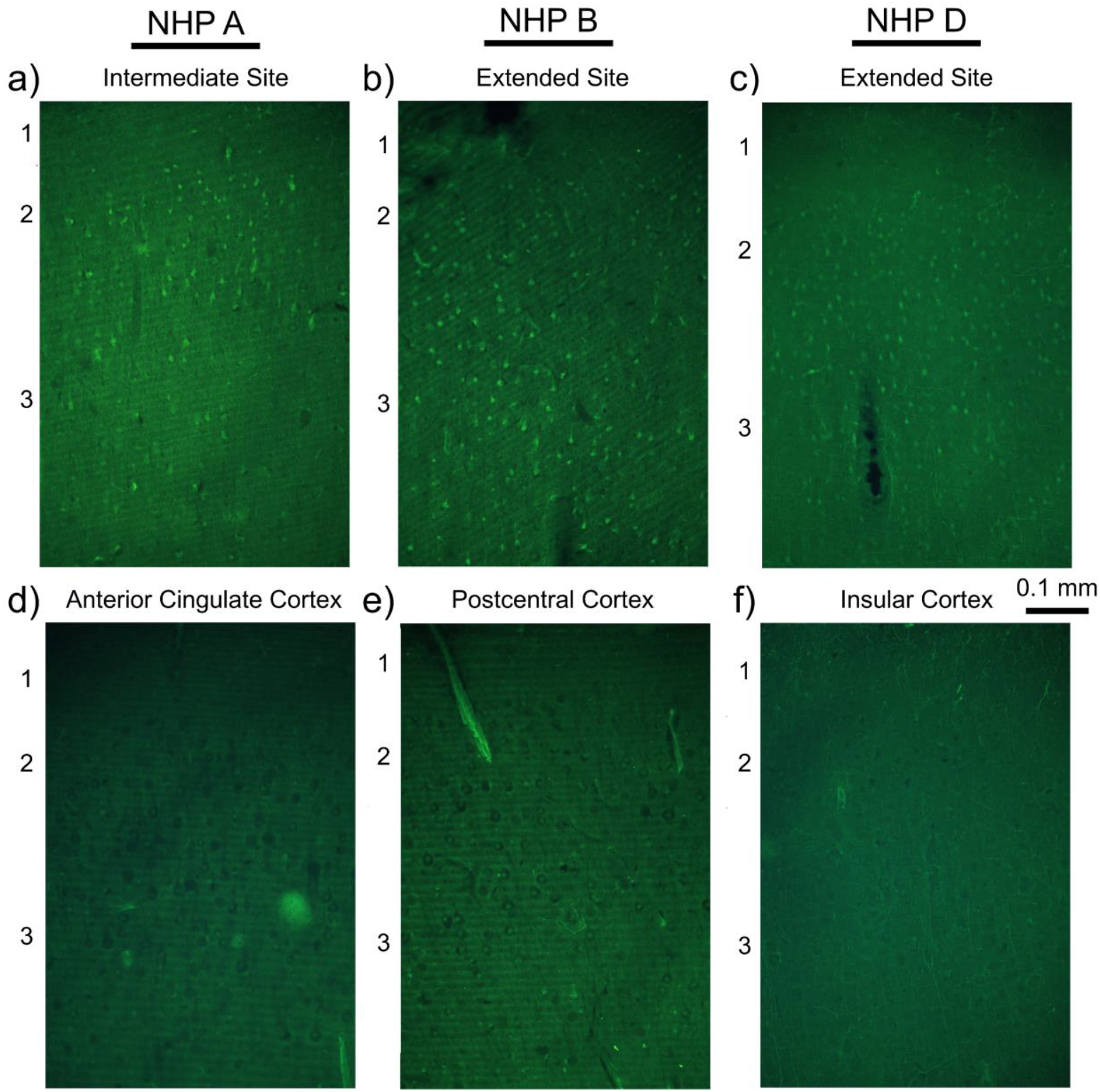
FJC labeling at stimulation and negative control sites. Panels show exemplary FJC-stained sections from histological analyses, with FJC-stained cells visible in bright green. Cortical layers are denoted on sides of panels, with common 0.1 mm scale bar provided. Each column of panels is from a single NHP. FJC-labeled cells are visible in regions associated with tFUS stimulation (a-c), including in the single-day intermediate site (a) and in extended-duration sites (b-c). FJC-labeled cells are not seen in negative control sites taken in the same subject far from the tFUS-targeted territory (d-f), supporting histological methodology.

**Figure 5.**
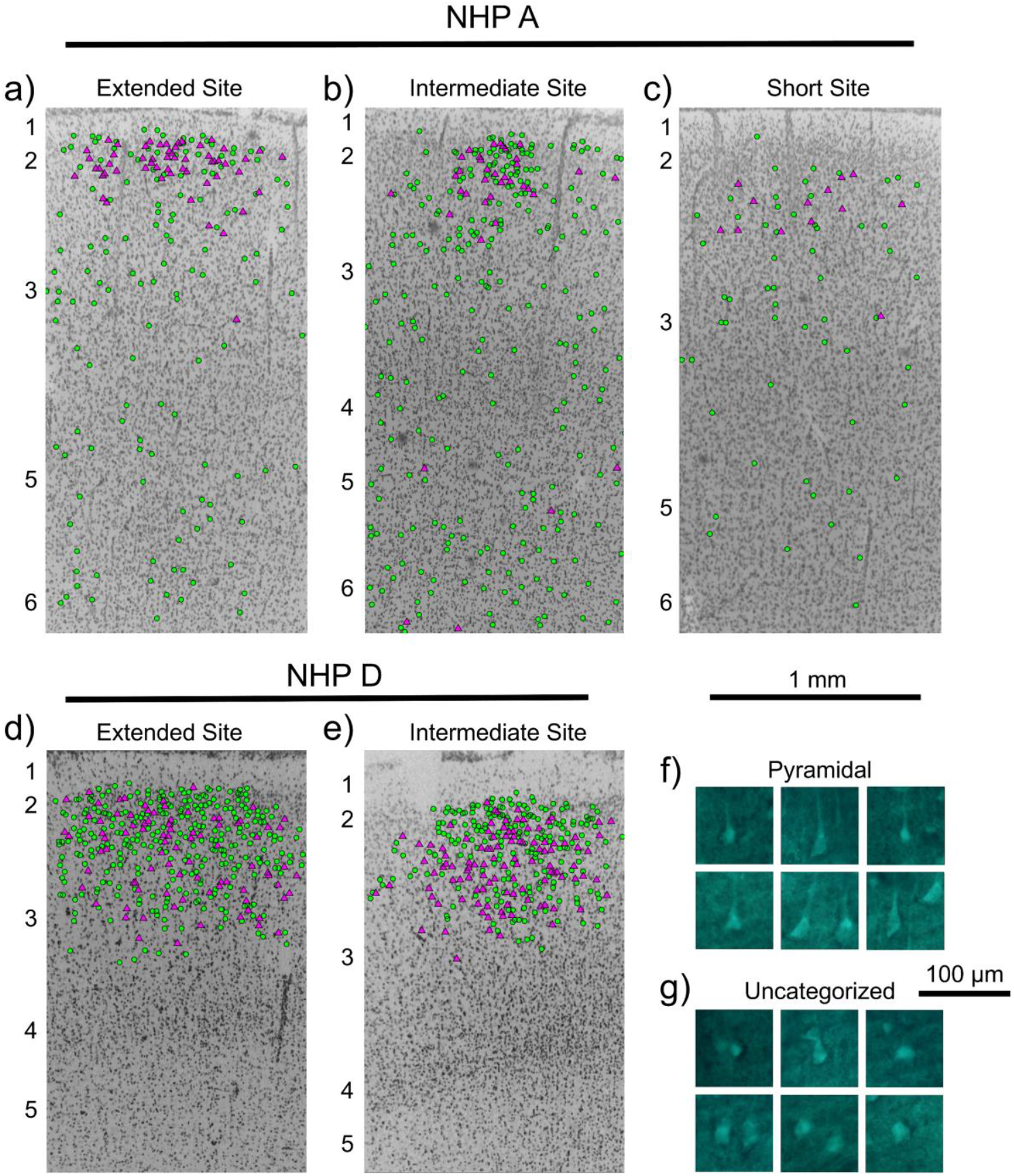
FJC labeling of pyramidal and uncategorized cells. (a-e) DAPI-stained sections of regions containing different stimulation sites in NHPs A and D; cell nuclei are stained gray, and FJC-labeled pyramidal and uncategorized cells are overlaid with magenta triangles and green circles, respectively. Cortical layers are denoted on sides of panels, with common 1 mm scale bar provided. FJC-labeled pyramidal cells are found in the upper layers in all 3 site types. Uncategorized cells are spread across multiple layers in NHP A, but not in NHP D. The density of labeled cells is similar across extended and intermediate-duration sites but is greatly reduced in the short-duration site. Virtually no labeled cells were observed at negative control sites [not shown]. (f-g) Example stainings of pyramidal (f) and uncategorized (g) cells, with common 100 μm scale bar. Pyramidal cells show triangular profiles with staining extended into the apical dendrite. Uncategorized cells are polygonal in shape and do not show staining in processes.

At extended-duration tFUS stimulated locations, FJC labeled cells were clustered in approximately 1.5 mm-diameter ovals around identified sites, with dense labeling within the central 1.0 mm region (Figure 5a-d). The intermediate-duration sites of NHPs C and D showed approximately 1 mm-diameter clusters of FJC-labeled pyramidal and uncategorized cells in layers 2-3 (Figure 5e). The 1-day intermediate-duration site in NHP A also showed a smaller-diameter, similarly dense cluster of FJC-labeled pyramidal cells in the upper cortical layers, though small numbers of labeled pyramidal cells were visible in deeper layers (Figure 5b). Additionally, FJC-labeled uncategorized cells were visible across multiple layers as in the 3-day extended-duration site. Short-duration sites in NHPs A and B showed sparse FJC-labeling. Again, FJC-labeled pyramidal cells were localized to layers 2-3, and uncategorized cells were visible across layers at a lower concentration than other sites (Figure 5c). Virtually no FJC-labeled cells were visible at deeper negative control sites, including the insular or cingulate regions lying directly under the tFUS-stimulated sites in areas 3a-4 (Figure 4d-f).

Table 2 lists the values of the MSD among labeled cells at each stimulated site, as well as the fraction of DAPI-labeled cells at each site that were also FJC labeled. According to this table and Table 1, both the extended-duration and the intermediate-duration sites in NHP A were tFUS stimulated at nearly the same intensity and showed comparable (statistically insignificant at p=0.26) MSD values and FJC-labeled DAPI fractions, despite different numbers of stimulation sessions. On the other hand, its short-duration site showed much lower labeling density (p<0.00001), despite being stimulated at higher tFUS intensities. In NHP B, FJC labeling density of its short-duration site was much lower than its extended-duration site (p<0.00001), despite being stimulated at much higher intensity. These outcomes suggest that the total amount of stimulation time a cortical site is subjected to in a single session is important for how much FJC labeling it will produce, while the number of such stimulation sessions less important.

**Table 2.** Stimulation site labeling. Number and percentage of and mean shortest distance (MSD) among FJC-labeled cells observed within a 1.25 x 1.25 mm^2^ region of interest (ROI) with densest FJC labeling at each noted site. MSD values are mean ± standard error of the mean; lower MSD values indicate greater density. Values for short-duration sites taken from individual site with densest labeling in given NHP (“Short (Max)”). Includes labeling information at strong positive control site (electrode damage to a blood vessel) and at clusters of FJC-labeled cells distant from tFUS focuses in NHPs B and D. Deep negative control sites had virtually no FJC-labeled cells.

| NHP | Site Type | FJC-Labeled Cells in ROI (#) | Fraction of DAPI-Labeled Cells with FJC Labeling (%) | Mean Shortest Distance MSD ($\mu\text{m}$) |
| --- | --- | --- | --- | --- |
| A | Extended | 153 | 11.2 | $40.6 \pm 2.1$ |
| | Intermediate | 197 | 11.4 | $36.5 \pm 1.7$ |
| | Short (Max) | 59 | 5.9 | $66.2 \pm 5.0$ |
|  | Deep | ~0 | ~0 | (~Inf) |
| B | Electrode Damage | 457 | 22.3 | $26.6 \pm 0.6$ |
| | Extended | 293 | 20.3 | $31.5 \pm 0.8$ |
| | Short (Max) | 91 | 5.0 | $50.4 \pm 2.8$ |
| | Distant | 67 | 7.3 | $56.1 \pm 3.8$ |
|  | Deep | ~0 | ~0 | (~Inf) |
| C | Extended | 130 | 17.9 | $27.4 \pm 1.0$ |
| | Intermediate | 259 | 27.2 | $20.0 \pm 0.5$ |
|  | Deep | ~0 | ~0 | (~Inf) |
| D | Extended | 384 | 20.2 | $24.8 \pm 0.5$ |
| | Intermediate | 315 | 18.8 | $24.2 \pm 0.5$ |
| | Distant | 306 | 23.8 | $24.8 \pm 0.5$ |
|  | Deep | ~0 | ~0 | (~Inf) |

In NHPs C and D, tFUS stimulation duration was made more than 3 times longer at the extended-duration sites than at the intermediate-duration sites, while stimulation intensity was reduced by about 50% (Table 1). As a result of this balancing of stimulation duration and intensity, FJC labeling densities at the extended- and intermediate-duration sites were similar (p=0.30) in NHP D, or even lower at the longer-duration site (p<0.00001) in NHP C.

Finally, the recording electrode inserted into NHP B pierced a blood vessel, producing tissue damage and prominent FJC labeling. Serving as a positive control, this labeling can be compared with that produced by tFUS stimulation. According to Table 2, the density of FJC labeling and its DAPI fraction at the extended-duration site were only slightly lower (p=0.031) than at the electrode damage site, making neurodegeneration caused by tFUS stimulation comparable to that caused by local blood vessel transection.

### FJC Labeling at Distant Sites

Sparse FJC labeling extended several millimeters away from the tFUS-stimulated cortical sites in all 4 NHPs. Just as in the stimulated sites, the scattered labeled cells were found in all cortical layers in NHPs A and B, but were confined to the upper layers in NHPs C and D. An example of such widely distributed FJC labeling, observed in NHP B, is shown in Figure 6. In contrast to this scattered labeling, not a single FJC-labeled cell was found in any of the 4 NHPs outside the sensorimotor cortical territory below the tFUS transducer array, including at deeper negative control sites (Figure 4d-f). This absence of any background FJC labeling suggests that the scattered FJC labeling outside tFUS-stimulated sites is related to that stimulation. In addition to broadly distributed FJC labeling, NHPs B and D each had a distinct cluster of FJC-labeled cells confined to the upper layers 4-5 mm anterior (NHP B; Figure 6a) or posterior (NHP D) of the tFUS stimulation site. In NHP B, the density of FJC labeling in its distant cluster was comparable to that at the short-duration stimulation site (Table 2; p=0.72). But in NHP D, the density of FJC labeling in its distant cluster was much stronger, comparable to that at its extended- and intermediate-duration stimulation sites (p=0.30 and p=0.44, respectively).

**Figure 6.**
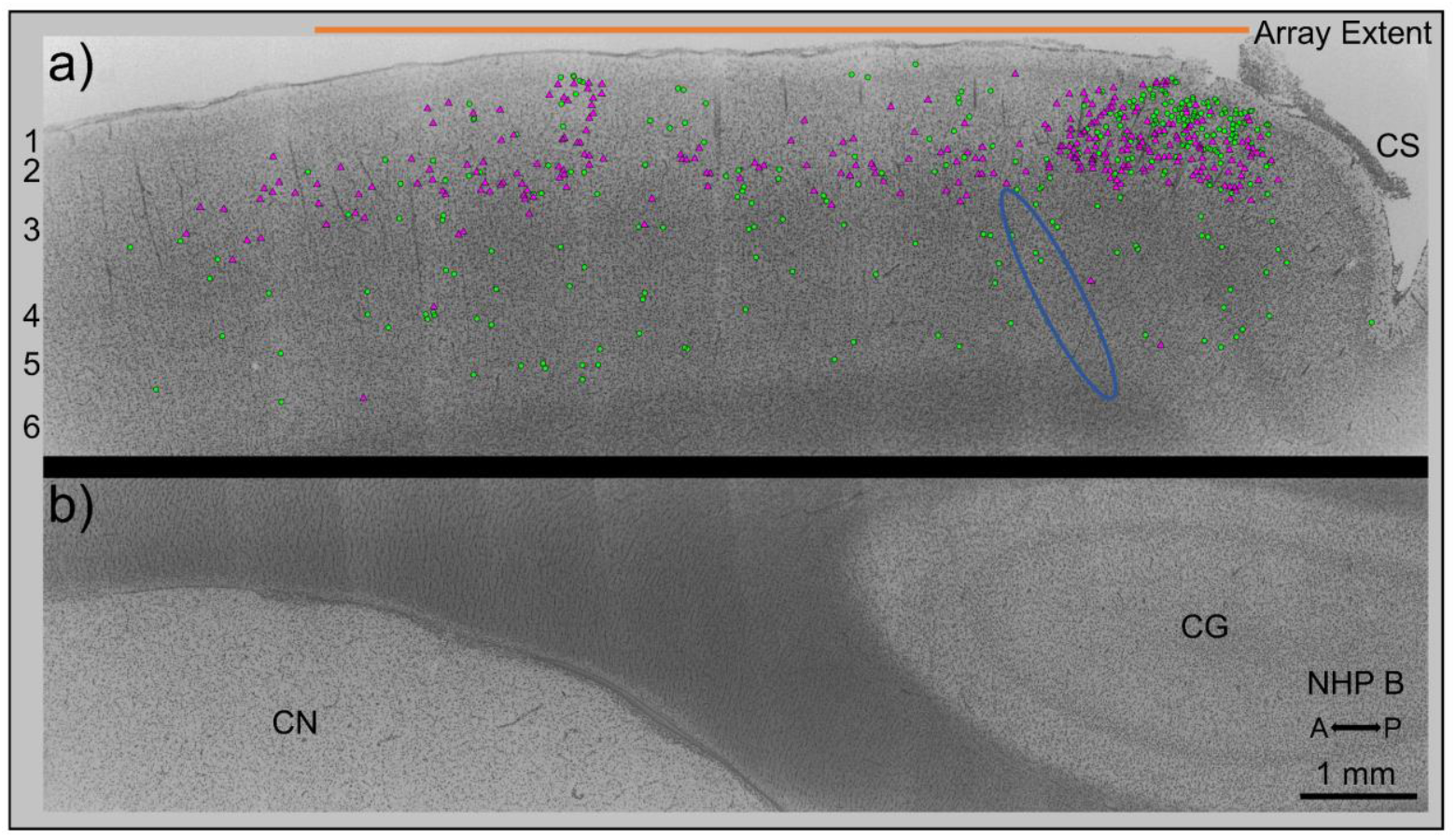
FJC labeling of targeted and distant sites. An exemplary DAPI-stained section from NHP B at the mediolateral location of the identified extended-duration site; cell nuclei are stained gray, and FJC-labeled pyramidal and uncategorized cells with are overlaid with magenta triangles and green circles, respectively. 1 mm scale bar and directional arrows (A = anterior, P = posterior) shown at bottom is common to both panels. (a) Section of crown cortex beneath tFUS array. Portion of section covered by tFUS array is shown as orange bar; actual array placed multiple mm above cortex. Reconstructed location of tFUS focus for extended-duration site shown as blue ellipse. Cortical layers listed left of panel, and CS is central sulcus. FJC-labeled cells are found densely clustered in the upper layers directly above (following radial lines) the ultrasound focus region. In addition, a scattering of FJC-labeled cells is found in all layers, as far as 8 mm anterior from the ultrasound focus region. The image shows the full 10 mm anterior-posterior spread of FJC-labeled cells. A distant cluster of FJC-labeled cells (center-left) is observed 5mm anterior from the focus region despite a lack of ultrasound foci in the vicinity. (b) Paired region of deep cortex 1 mm directly beneath stimulated crown cortex in top panel, serving as negative control. CN is caudate nucleus, and CG is cingulate gyrus. Virtually no labeled cells were observed in such regions.

### Behavioral Responses

Figure 7 plots the task performance for each NHP over the course of the study. Performance on the task trended positively over time for each NHP, consistent with task learning. Task performance did not degrade in any NHP following tFUS stimulation.

**Figure 7.**
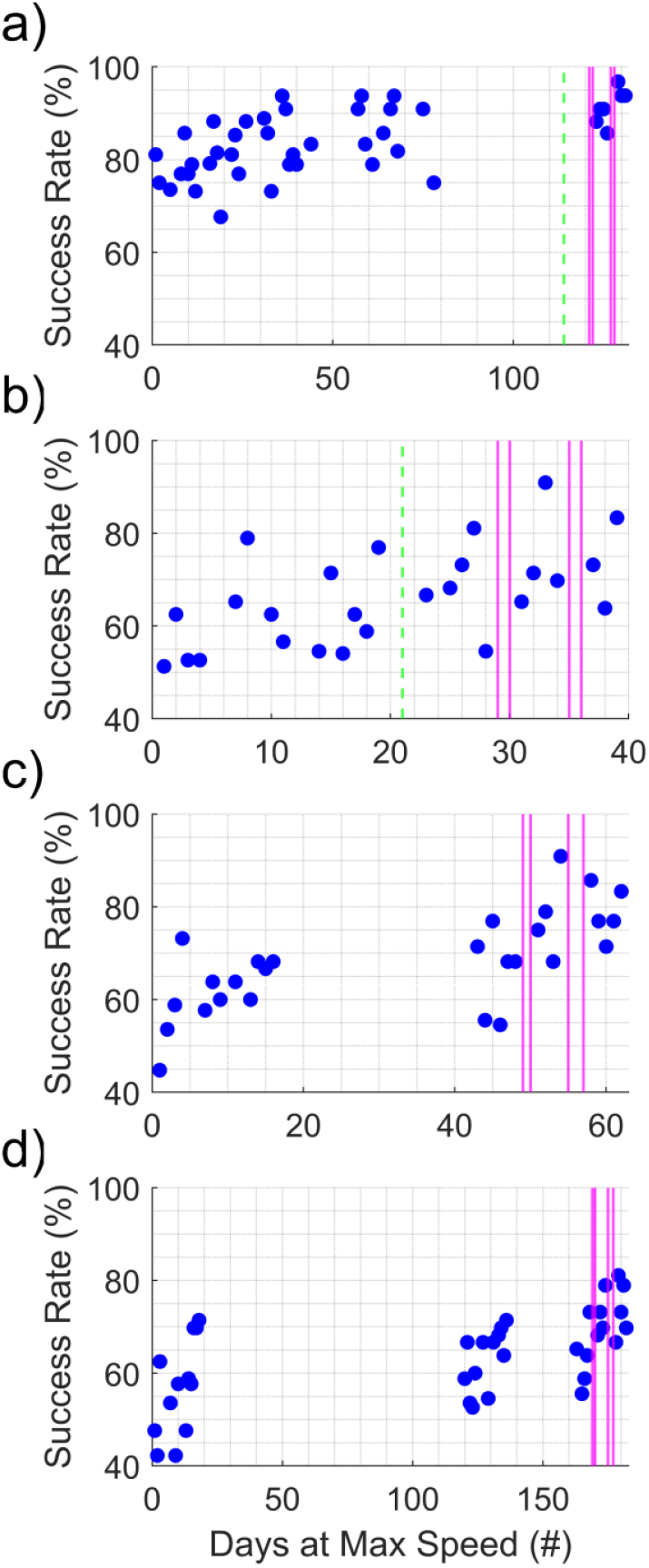
Behavioral testing before and after stimulation. Success rates at pellet-picking task in NHPs A-D in corresponding panels (a-d). Dotted green lines indicate time of electrode implantation in NHPs A and B. Magenta lines indicate stimulation sessions for each NHP. No NHP showed degradation on the performance task over the course of testing or following stimulation.

### Effects of Stimulus Parameters

Next, we related tFUS stimulus parameters to the degree of neurodegeneration seen at each stimulation site. Figure 8 plots the stimulus intensity and FJC labeling density (MSD) against *T*_*E*_ at each site. All examined tFUS-related sites displayed FJC-labeling despite a broad range of applied intensities, though MSD values unexpectedly increased with increased intensity, reflecting reduced labeling densities. In contrast, MSD values trended downwards with greater *T*_*E*_, reflecting greater labeling densities. MSD was similar for sites with *T*_*E*_ greater than 20 sec, suggesting a ceiling effect. This ceiling closely resembled the labeling density of the positive control electrode damage site in NHP B.

**Figure 8.**
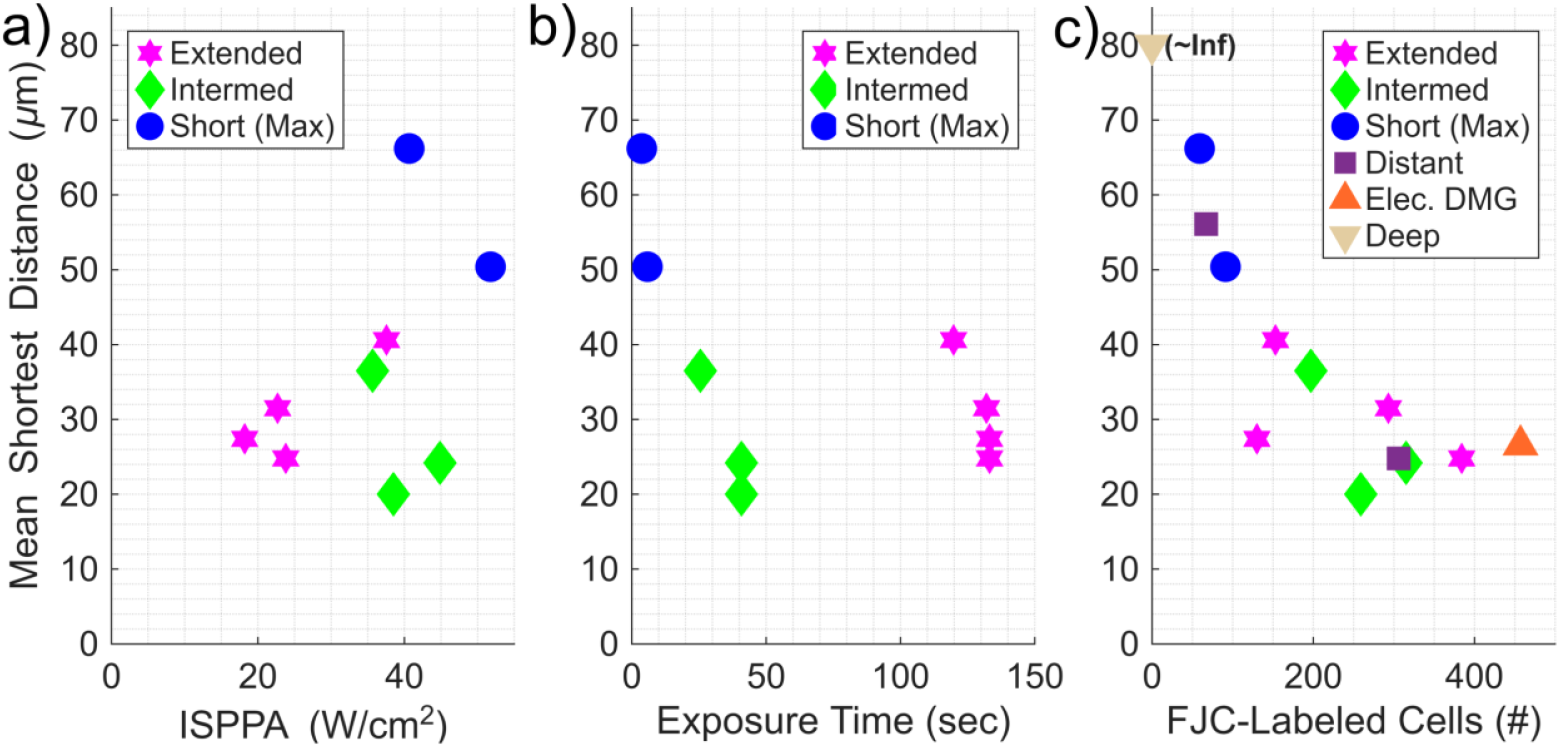
Stimulation site labeling and trends. Mean shortest distance (MSD) values among FJC-labeled cells at each measured site across all NHPs, with site types denoted by color/shape. Values for short-duration sites taken from individual site with densest labeling in given NHP (“Short (Max)”), assuming an average ISPPA and *T*_*E*_ from short-duration sites in given NHP. (a) Trends in MSD based on site intensity. Sites with higher ISPPA values have higher MSD values, reflecting reduced labeling density. (b) Trends in MSD based on site exposure time. Short-duration sites have large MSD values compared to both intermediate and extended-duration sites, which are highly similar. (c) Trends in MSD with number of labeled cells in each observed site ROI. MSD values for short-duration sites cluster away from other-duration sites, which cluster together and with an electrode-damage site. Distant sites have similar MSDs and FJC-counts to tFUS sites. Deep negative control sites had virtually no FJC-labeled cells.

## Discussion

### Neurodegeneration at Target Sites

Our histological analyses indicate that tFUS stimulation can result in neurodegeneration in certain stimulus parameter regimes. FJC labeling consistently showed neurodegeneration in proximity to stimulation sites in all NHPs, and the MSD of labeling decreased with greater *T*_*E*_, reflecting increased density. Clusters of FJC-labeled cells at extended-duration and intermediate-duration sites showed similar MSDs, suggesting the varied stimulus patterns had a similar and likely-saturated net effect on cortical tissue. In contrast, the greatly reduced FJC-labeling at short-duration sites despite their higher intensities suggests that the neurodegeneration was at least partly driven by *T*_*E*_.

Some differences among NHPs and deviations from simulated results can be attributed to the experimental setup. The lack of FJC-labeled cells in the deep cortical layers in NHPs C and D could due to a lack of surgery-related damage or a difference in labeling effectiveness based on sample acquisition dates [34, 35]. The central regions of clusters were larger than predicted, though this could be attributed to minor variations in transducer placement across multiple sessions. Additionally, the wider spread of labeling for extended-duration sites is likely due to their comparatively transverse focus orientation (see Figure 6). However, the presence of any neurodegeneration represents an unexpected result in this initial study.

### Neurodegeneration at Distant Sites

The appearance of scattered FJC labeling outside the directly stimulated cortical sites, extending as far away as 8 mm in the anterior-posterior direction, as well as denser clusters of labeled cells 4-5 mm away from the stimulated sites is especially surprising. Such labeling indicates that focused stimulation is capable of inducing neurodegeneration at sites far outside the focal region under the right conditions. The density of degrading neurons in observed clusters varied across NHPs but was consistent with other tFUS-associated sites, suggesting that neurodegeneration near and far from the focused site resulted from a common biological source. This is further supported by the neuroanatomical connections between distant sites and the primary stimulation targets.

### Behavioral Effects

Despite observed neurodegeneration, none of the NHPs were negatively impacted on the behavioral task. One interpretation of these results is that the levels of neurodegeneration seen here were not enough to affect system functionality. This could be due to the limited volumes within which degeneration occurred, where the number of altered cells was insufficient to affect manual dexterity or muscular function. Other cortical regions with overlapping receptive fields [36] could also be compensating for losses. The similarities between the 1-day and 3-day sites in NHP A suggest that additional tFUS sessions would not lead to increased neurodegeneration, thus maintaining overall function, but additional studies are warranted to confirm given the observed trends of degeneration with increased *T*_*E*_.

### Biological Sources for Neurodegeneration

tFUS-associated neurodegeneration observed in cortical tissues appears to be dependent on the cumulative duration and intensity of exposure near to or within targeted regions. It is unlikely neurodegeneration was caused by thermal effects as the maximum temperature increase predicted in this study in cortical tissue was 0.13°C. In simulations, it has been shown [37] that 1.9 MHz stimulation with extended *PTDs* [38] can significantly heat up tissue adjacent to the skull through diffusion, but as we used much lower intensities and much shorter *PTDs*, we would expect significantly less heating from such mechanisms during a single pulse train. Furthermore, temperatures would be unlikely to accumulate over pulse train repetitions given our extended 10 sec *ISI*.

It is not clear if *T*_*E*_ alone is sufficient to cause neurodegeneration, or if a minimal stimulation intensity may be required to meet a threshold of exposure. That said, it is also unlikely that tFUS intensity alone drove the types of neurodegeneration seen here. The maximum *MI* experienced given our *f*_*0*_ was less than the FDA limit of 1.9, suggesting the effect was not due to cavitation, and our *ISPPA* for all sites were below FDA limit of 190 W/cm^2^ for mechanical damage [16]. Comparatively low levels of FJC-labeling at sites with the highest-intensity foci and the appearance of labeled cell clusters far from tFUS focal regions indicate the effect is not solely due to the intensity of tFUS. The lack of high-intensity side lobes in simulations reduces the likelihood such clusters were due to off-site focusing, though the localization of scattered FJC labeling to the region directly under the CMUT array does suggest effects were nonetheless tFUS-related.

A recent work [39] highlighted strain resulting from the acoustic radiation force (ARF) as one means by which tFUS impacts neural tissue. ARF-strain is reportedly predominantly shear, and using estimation methods discussed in that article, it scales linearly with intensity but at least quadratically with *f*_*0*_. While prior safety studies in NHPs have used intensities in a similar range to ours [40], the much higher *f*_*0*_ used here likely induced ARF-strain levels 20-80x greater (see Appendix A), potentially leading to neurodegeneration. The resulting effects would likely build up over time with repeated stimulus applications, as neuron membranes can be compromised under complex mechanical deformation [41].

However, the appearance of scattered and clustered neurodegeneration far from tFUS foci suggests the distant neurodegeneration is underlaid by tFUS’ impact on axons rather than on somata themselves. Our stimulation foci were biased towards the middle-deep layers and white matter below to avoid inadvertently focusing in and heating the skull, and degenerating neurons were localized largely in superficial cortical layers whose small-diameter descending axons ran through the focused regions. The appearance of FJC-labeled somata may have resulted from retrograde degeneration induced by axon damage [42, 43]. Alternately, axon damage [44] or excessive tFUS-induced neuron firing [45] could have produced excitotoxic conditions [46] in downstream regions. A recent tFUS study [5] did not observe any negative behavioral effects when targeting white matter in humans, though they utilized sub-MHz *f*_*0*_ and comparatively-low DC stimuli.

The location of the distant clusters of FJC-labeled cells noted in NHPs B and D supports the connectivity theory. These distant clusters were located in Brodmann’s areas 6 and 2, respectively, which are anatomically connected to the primary stimulation sites for both NHPs, in the posterior part of area 4 and anterior area 3a [47-50]. Furthermore, the columnar (radial) trajectory of the axons of the FJC-labeled cluster in NHP B, associated with extended-duration stimulation (Figure 6, right), aligned with the projected focus of tFUS stimulation. We posit the sparse-but-widespread FJC labeling is due to local connections among stimulated sites. Since tFUS stimulation of white matter likely affects regions beyond the initial targets [51], this strongly indicates that stimulation targeting and protocols must be carefully considered to avoid undesirable consequences.

## Conclusion

Extended tFUS exposure and subsequent histological analyses revealed unexpected neurodegeneration in an initial study testing 4 squirrel monkeys. The degeneration was related to the total exposure time experienced by a given focus site, which may have been partially driven by the high operating frequency of the stimulus. However, it is likely that the degeneration was also related to the stimulation of axons in the region, which also induced neurodegeneration at sites distant from the chosen target. A clearer understanding of how repeated tFUS affects targeted tissues and compositions is needed to better capture all the appropriate biological factors that contribute to neurodegeneration.

While it remains uncertain if the effects shown here are typical of tFUS stimulation, the results of this study suggest further examinations are warranted. Before transcranial focused ultrasound stimulation can be applied in a long-term capacity, key features of safe and effective stimulation paradigms need to be identified and examined, which will require additional investigations into the effects of extended tFUS exposure on different cortical regions or cytoarchitectures.

### Credit Authorship Contribution Statement

Andrew Brna assisted in NHP stimulation session design, led tFUS NHP stimulation sessions, aided in result interpretation, and authored the initial text and figures. Oleg Favorov conceptualized and participated in NHP stimulation sessions, performed surgeries, ran electrode recording equipment, prepared and interpreted histological slides, and edited text and figures. Timothy Challener ran NHP behavioral testing and assisted in NHP experiments. Muhammetgeldi Annayev designed, fabricated, and tested the tFUS arrays. Ali Biliroglu integrated the electronics and arrays in the stimulator probe and operated the device in NHP and hydrophone experiments. Erkan Kemal operated the ultrasonic stimulator in NHP experiments. Yalcin Yamaner and Omer Oralkan conceptualized the tFUS system and oversaw its design, integration, and testing. Derek Eidum performed Fullwave simulations, derived NHP phase values, and assisted in stimulation session design. Stephen Simons and Michael Weisend aided in planning, execution, analysis, and interpretation of NHP experiments. Patrick Connolly led the research project as principal investigator, conceptualized the tFUS system and NHP experimental design, oversaw experimental efforts, interpreted results, and edited text and figures.

## Supporting information

High_Res_Histology_Figures

## Acknowledgements

Histological services were provided by the Histology Research Core Facility in the Department of Cell Biology and Physiology at the University of North Carolina, Chapel Hill.

The authors gratefully acknowledge the expert veterinary assistance of Steven T. Shipley, Nicole Massoud, and Leslie Whitfield of UNC Division of Comparative Medicine.

The authors additionally thank Sabrina Brna for providing artistic renditions of NHPs for article figures.

This work was supported by the Naval Information Warfare Center (NIWC) and the Defense Advanced Research Projects Agency (DARPA) under Contract No. N65236-19-C-8013. The views, opinions, and/or findings contained in this paper are those of the authors and should not be interpreted as representing the official views or policies, either expressed or implied, of the NIWC Atlantic, DARPA, or the Department of Defense.

Distribution Statement “A” (Approved for Public Release, Distribution Unlimited)

## Appendix A: Acoustic Radiation Force Strain Estimates

Nandi et al. [39] describe a method of estimating strain resulting from acoustic radiation force (ARF). The method consists of “normalizing the displacement [resulting from ARF] to the wavelength [of the ultrasound beam]”. We summarize this method as Equation S1.

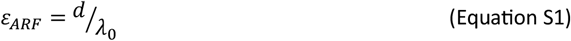

where ε_ARF_ is the ARF-induced strain, *d* is ARF-induced tissue displacement (m) and *λ*_*0*_ is the wavelength of the ultrasound beam (m). In neural tissue, *λ*_*0*_ is related to the operating frequency *f*_*0*_ by Equation S2.

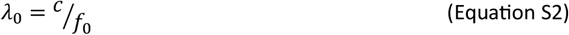

where *c* is the speed of sound in neural tissue (1468.4 m/s) and *f*_*0*_ is the operating frequency (Hz). In an earlier work, Ye et al. [27] provide a calculation for ARF-induced tissue displacement which assumes displacement is proportional to ARF by some constant; we provide their equation as Equation S3.

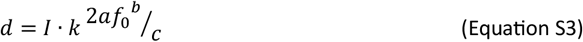

where *I* is the spatially varying pulse average intensity experienced at the focus site (W/m^2^), *f*_*0*_ is the operating frequency (MHz), *a* is the absorption coefficient of cortical tissue (2.4 Np/m/MHz^-b^), *b* is the power law coefficient (1.18), *c* is the speed of sound in neural tissue (1468.4 m/s), and *k* is some tissue-specific constant. Combining S1-S3 and reconciling known units, we estimate ε_ARF_ as Equation S4, which is consistent with Nandi et al.’s statements that the value is proportional to both I and f_0_ squared (if not more so) as *b* is greater than 1.

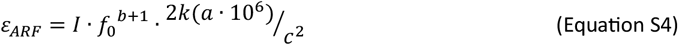

While Ye et al. do not provide a value for *k*, Equation S4 is still useful to make comparisons of *ε*_*ARF*_ among works operating on similar tissues. Taking *ISPPA* as a maximum value for the pulse averaged intensity experienced during stimulation, we calculate values of 1.46x10^6^*k* and 5.24x10^6^*k* for as the range of *ε*_*ARF*_ experienced across our stimulated sites using our minimum and maximum calculated derated *ISPPAs*.

Gaur et al. [40] ran a similar study in macaques and sheep looking at the safety of extended tFUS stimulation. This study applied a 270 kHz *f*_*0*_ stimulus with a maximum *ISPPA* of 51.6 W/cm^2^ in a macaque over 500 pulse trains, yet they saw no damage during histological analyses. Using Equation S4 and the same constants used in our study, we estimate such stimuli to experience *ε*_*ARF*_ of 6.62x10^4^*k*. While their *ISPPA* value was within the range tested in our study, the calculated minimum and maximum *ε*_*ARF*_ for our study were 22.1x and 79.3x (or approximately 20-80x) the value of Gaur et al. due to the difference in *f*_*0*_ values. This supports the theory that *f*_*0*_ is likely involved in the neurodegeneration seen in our study, though our other results such as those in Figure 8 also suggest duration is involved alongside *f*_*0*_.

