## Supplementary figures and images for "Neurodegeneration Associated with Repeated High-Frequency Transcranial Focused Ultrasound"

### Figure_5.png

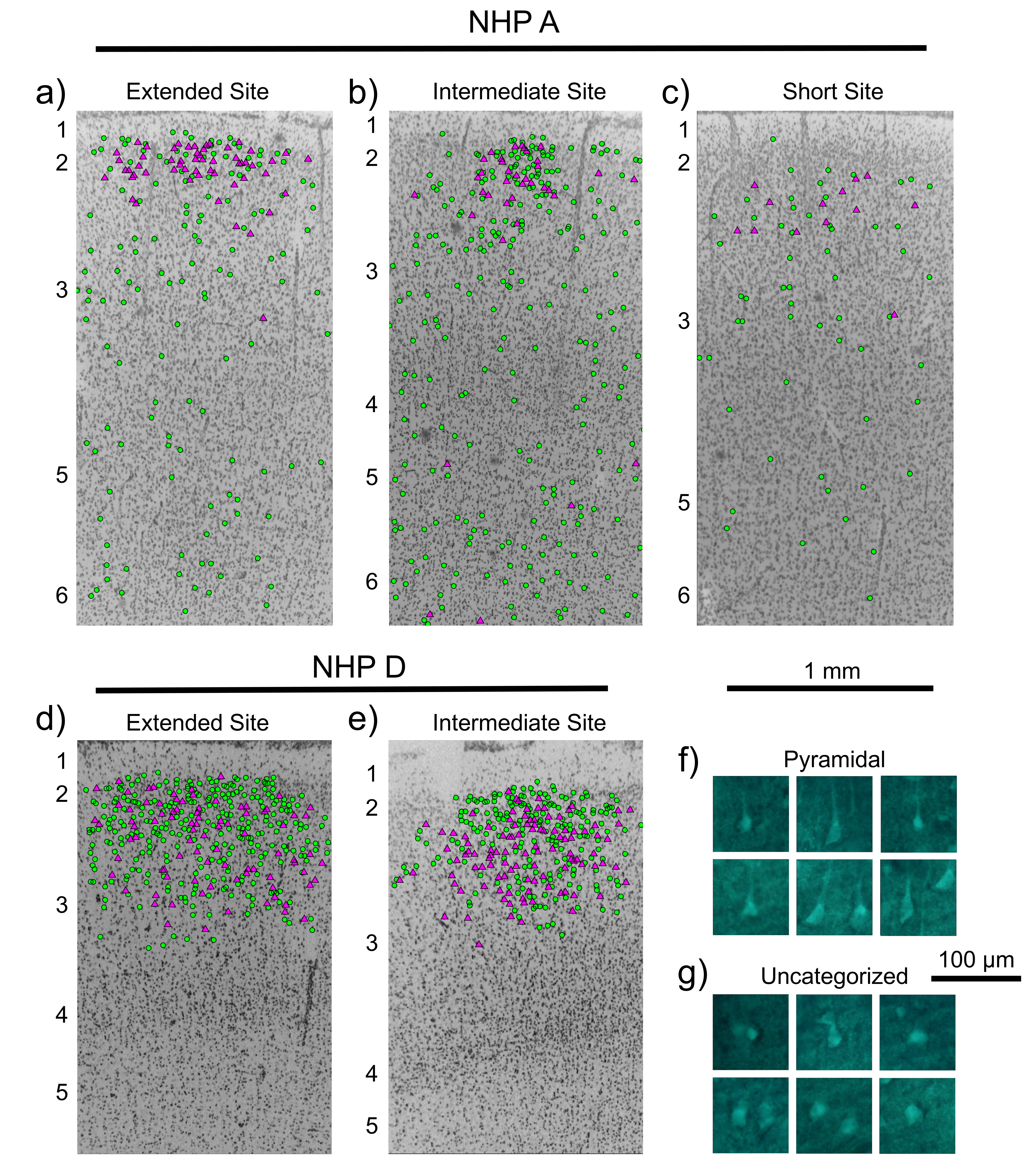
